# The gravitation-driven stress-reduced urothelial barrier in toad bladder urothelium

**DOI:** 10.1101/2021.06.20.449200

**Authors:** Tan Xinli, Wang Danmei, Fan Shouyan, Xu Xiaochi, Guo Hui, Emmanuel Kwao Teye, Merveille Nnanga Ndjike Michele, Wang Yang

## Abstract

The urinary bladder urothelial are highly specialized epithelia that protect the underlying tissues from mechanical stress and seal them from the overlying fluid space. To better understand the maintaining permeability induced electrical potential roles played by urothelial in the bladder, we established a protocol of gravitation stress in toad urothelial, observed the transmembrane potential difference variation.

**Method:** The toad urothelial were mounted in a using chamber which the chamber was separated to two solution spaces, and stable with 0.9% saline solution. The electrodes were settled on the surface of each side of the preparation, serosal side definite as cathode. The using chamber was settled in the centrifugal rotor and under 300 rpm rotation to obtain a vertically +4G gravitation on serosal chamber 5min.

**Result:** a transient transmembrane potential difference increasing was observed after adding CaCl_2_ (3% solution) in serosal chamber. The amplitude increasing phase included a rapid and a slowly ascending phase. In gravitation stressed urothelial preparation, CaCl_2_ induced transient phase was significantly increased, furthermore the secondary slowly ascending phase was much more amplified on its amplitude axis and significantly prolonged on the time scale than that evoked in control preparations. The evoked total amplitude increasing were 10 times higher than that in control.

**Conclusion:** The urinary bladder epithelial layer has a structure which regulates ion permeability as a barrier. The tight junction plays an important role as the intercellular coupling in the apical side of the epithelial cell. On the other hand, it is known that the ion channel exists on the epithelial cell membrane and regulates the physiological process. The gravitation stress weakened the tight junction. The transmembrane potential difference was enhanced both on its amplitude and prolonged time. The gravitation stress induced hyperpolarization that evoked by CaCl_2_ is one kind of Cl^-^ transfer from serosal chamber in which high Ca^2+^ in the urothelial basal membrane activated the calcium-activated chloride channels. This outwardly rectifying chloride channel induced hyperpolarization can be blocked by *Nppb*.

## Introduction

The urothelial lines the inner surface of the urinary bladder, where it forms a tight barrier that allows for retention of urine, while preventing the unregulated movement of ions, solutes, and toxic metabolites across the epithelial barrier. The urothelial permeability barrier is made up of three components: apical membrane, tight junctions, and the trafficking mechanism that inserts and removes apical membrane in response to filling and emptying of the bladder. The apical membrane contains specialized lipid molecules and uroplakin proteins that appear to function in concert to reduce permeability of the apical membrane to small molecules such as water, ammonia, protons, and urea ^[*1*]^. The tight junctions sharply restrict the movement of ions, such as Na^+^, K^+^, and Cl^-[*2*]^. The permeability barrier must be maintained even as the organ undergoes cyclical changes in pressure as it fills and empties. Beyond furthering the understanding of pressure induced barrier function, new analysis of the urothelial is providing some updated information about the plaques (detergent-insoluble membrane/protein domains) are formed at the apical plasma membrane of the surface umbrella cells ^[*3*]^. The mechanical stimuli such as pressure alter urothelial exocytic and endocytic traffic ^[*4*]^. The urothelial barrier to ion, solute, and toxin flux must also adapt to large variations in pressure as the bladder fills. Defects in plaque assembly increase membrane permeability and may lead to diseases such as vesicoureteral reflux ^[*5, 6*]^. Therefore, urothelial studies provided the clues of its sense to mechanical stimuli such as pressure, and transduce changes in these stimuli into cellular events such as membrane traffic. The studies indicated that increased pressure stimulates exocytosis and endocytosis in umbrella cells and modulated by purinergic signaling cascades, Ca2þ, and cAMP ^[*7*]^. Nothing is known about regulation of pressure induced cross barrier electrical potential differences, however this is important for understanding its physiological excitation. Future analysis of this pathway may lead to a better understanding of mechanical stress-independent pathways of internalization. In an Ussing chamber model, it is possible to analyses the urothelial layer potential difference *in vitro*. Some reports suggested that protamine irrigating increased the barrier permeability to urea, a 3.5% urea movement across the membrane, appeared that the impairing of urothelial surface may increasing barrier in modulating both charged and uncharged small molecule movement ^[*8*]^. In human being, the urothelial barrier deficits, suburothelial inflammation and sensory proteins expressed in the bladder mucosa usually combined with detrusor underactivity. The expression of zona occuldens-1, E-cadherin, tryptase and apoptosis levels in the suburothelium, β3-adrenoceptor, M2 and M3 muscarinic receptors, P2X3 receptor, and inducible and endothelial nitric oxide synthase were involed. The lower E-cadherin expression, lower expression of M2 and M3 muscarinic receptors, was detected in these patients ^[*9*]^. Urothelial dysfunction altered sensory protein expressions, prominent the impaired urothelial signaling and sensory transduction pathways ^[*10*]^. With the availability of Ussing chamber model, we can explore the reason of ionic permeability that regulate the transmembrane potential difference. Some studies demonstrate that using chamber method is help to understand the urothelial undergoes characteristic function and structural response to injury, provide an approach to defining how this process is regulated. The understanding of this mechanism is able to acquire how the urothelial repairs itself may lead to important insights into the care of the human being with cystitis and may also shed light on the mechanisms by which bladder cancer develops ^[*11*]^. The environmental differences between the apical and basal surfaces of the urothelial, such as osmotic pressure and hydrostatic pressure influence the tight junction related barrier action. Especially, the complex fiber protein network is constructed in the epithelial cell junction maintained the low permeability. It was clarified that the mechanical stretch reduced 10 times of the resistance of the transcranial electrical resistance (TER) and the umbrella cell junction resistance ^[*12*]^. The membrane permeability and the membrane bound were changed by ion permeabilization. In this study, we investigated the effect of gravity acceleration on the ion permeability and excitability of the bladder epithelium.

## Methods

The toad (4 weeks, n = 10) was purchased from the experimental animal center of Hainan medical university The study protocol was approved by the ethics committee of Hainan medical university. All animal experiments were performed in accordance with the guidelines of the animal care and use committee of Hainan medical university. The toads sacrificed by foramen magnum pithing. After midline incision from the sciatic to the hypodermic center, the bladder body was separated by a glass probe. The gap with the total urinary bladder body was removed, then stable in Ringer’s solution for 10 min, room temperature. The urinary bladder urothelial wall was separated to two parts by ophthalmic scissors. One part was for control, while another part was for gravitation stress test. The urothelial layer was mounted on the Ussing chamber, therefore the Ussing chamber was consist of two chambers, apical membrane chamber (apical chamber) and basement membrane chamber (serosal chamber). The chambers were fulfilled with 0.9% sodium chloride solution and stabilized for 5 min, room temperature (**Figure 1**). The Ussing chamber was vertically centrifuged at 300 rpm 5min to obtain a 4G acceleration on its basement membrane. This is the gravitation stress induced mechanical shed urothelial preparation. After that the apical membrane and basement membrane was settled with electrode, the serosal side definite as cathode. The electrodes were connected to the polygraph device for recording the transmembrane potential difference (TPD). The urothelial membrane was activated by 3% CaCl_2_ solution on its serosal side. The enhanced potential amplitude and prolong time were analyzed. In a compared study the mechanical shed urothelial layer, the chemical shed with glycerin was prepared. For investigating the Ca^2+^ induced Cl^-^ movement crossing the urothelial layer, the chloride channel blocker *Nppb* was used for observing calcium-sensitive chloride currents.

**Figure 1.**
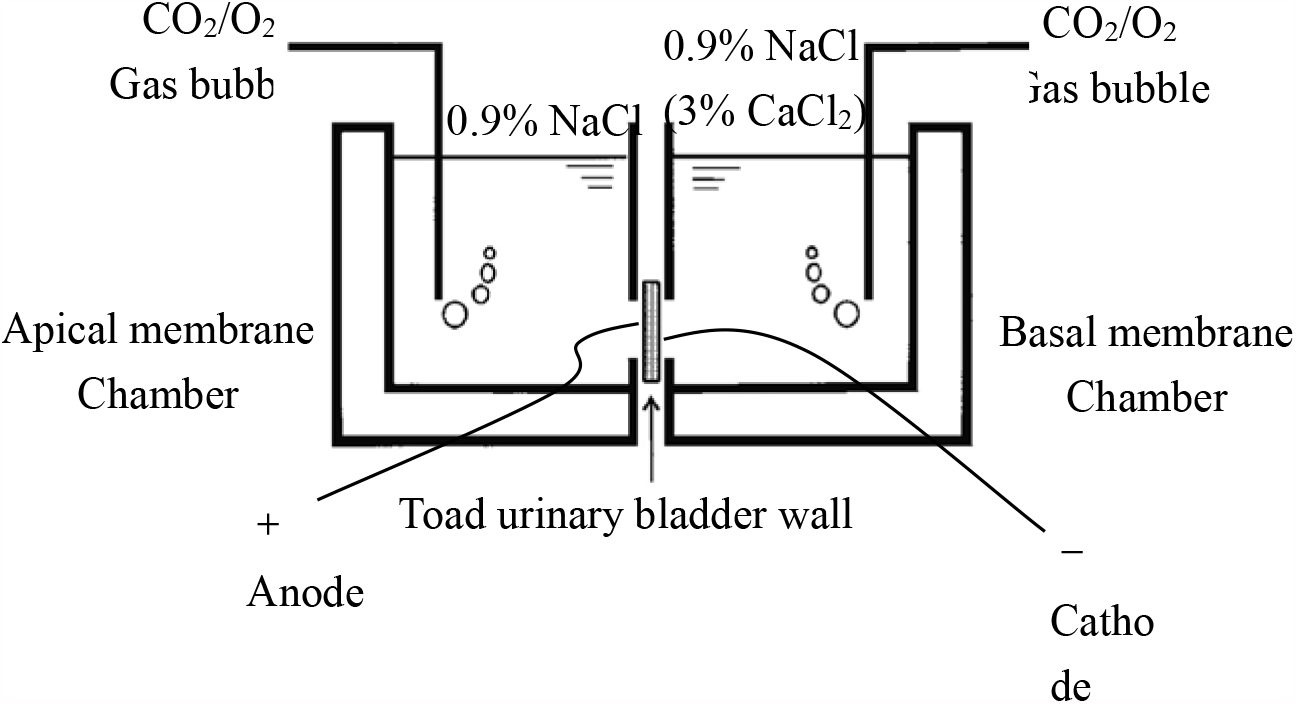
Ussing chamber for toad urothelial layer for transmembrane potential difference testing.

## Results

In control intact urothelial layer, CaCl_2_ activation induced an increasing amplitude of transient transmembrane potential difference (TPD). The amplitude increased rising phase included a rapid and a slowly ascending phase (**Figure 2a**, *). After the transient increasing phase, the amplitude maintained for a long period (arrow mark).

**Figure 2.**
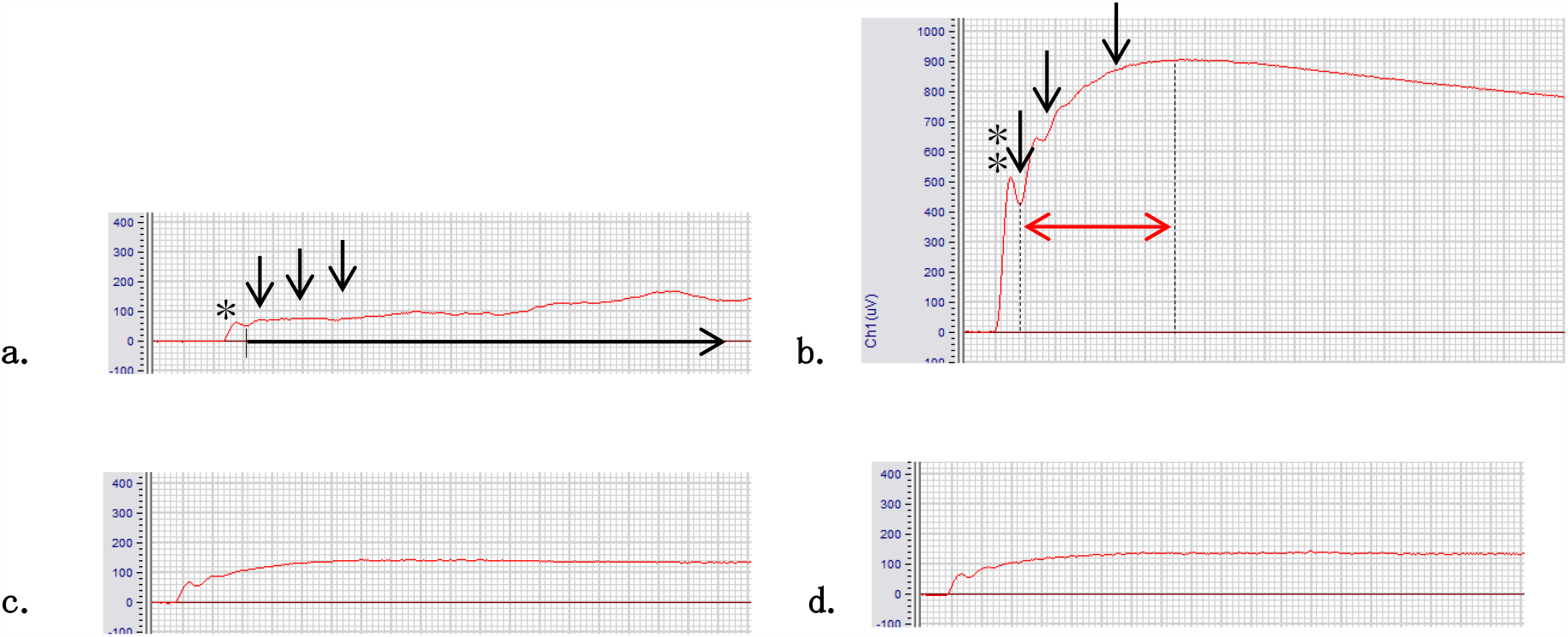
Transmembrane potential differences in prepared urothelial layer. a. Control intact preparation; b. Gravitation stress shed preparation; c. Glycerin skinned preparation; d. *Nppb* treated mechanical shed preparation

In gravitation stress induced mechanical shed urothelial preparation, CaCl_2_ induced transient rising phase was significantly enhanced, furthermore the secondary slowly ascending phase was much more amplified on its amplitude axis, and the duration was significantly prolonged on the time scale than the control intact urothelial preparation. The CaCl_2_ evoked transient rising phase was 10 times enhanced than that in control preparation (**Figure 2b**, **). The arrow mark indicated a typical secondary increasing after the transient rising phase. This secondary increasing continuously 6 times prolonged than the initial amplitude increasing (red arrow period). The urothelial layer was largely hyperpolarized. **Figure 2c** shown the CaCl_2_ activated TPD in glycerin shed urothelial preparation. The chemical shed did not obtain the significant increasing of CaCl_2_ activated TPD. **Figure 2d** was the *Nppb* intervened TPD in mechanical shed urothelial preparation. The TPD were significantly reduced because of blocking the chloride channel which indicated that the main compound of the CaCl_2_ activated amplitude increasing was Cl^-^. This further revealed that the Cl^-^ current induced the hyperpolarization in mechanical shed urothelial preparation.

## Conclusion

The bladder epithelial layer has a structure which regulates ion permeability as a barrier, and tight binding plays an important role as the intercellular coupling in the apical side of the epithelial cell. On the other hand, it is known that the ion channel exists on the epithelial cell membrane and regulates the physiological process. It was reported that the transurothelial electrical resistance decreased with the urinary tract injury using protamine sulfate ^[*13*]^. The mechanical stretch method is noticed as a method to weaken the tight coupling structure by minimizing the cell membrane damage unlike the chemical damage. Using this method, the gravitational stress (4G) was added to the basal membrane. The apical membrane and basal membrane permeability was deeply influence.

In control intact urothelial preparation, CaCl_2_ activated the Ca^2+^-activated chloride channels (CaCCs), induced a amplitude increasing of the TPD. This increasing make the urothelial to be in a slightly hyperpolarization on its apical membrane, therefore protect the apical membrane from the over excitation. The experiment results indicated that in the mechanical shed urothelial preparation, this hyperpolarization was drastic enhanced. The membrane excitation was inhibited by the significant Cl^-^ movement from the basal membrane toward to the apical membrane. In contrast, there was no significant increasing of such inhibition in chemical shed urothelial preparation.

The Using chamber using for investigating the mechanical shed urothelial permeability is a new challenged laboratory protocol for study the urothelial physiological excitation during chemical compound intervened. This will help to develop the new drug absorption system to improve the urinary bladder osmotic therapy.

## Acknowledgements

This research was carried out by the University Student Innovation Challenge promotion project (S202011810032).

